# Principal-stretch-based constitutive neural networks autonomously discover a subclass of Ogden models for human brain tissue

**DOI:** 10.1101/2023.01.14.524079

**Authors:** Sarah R. St. Pierre, Kevin Linka, Ellen Kuhl

## Abstract

The soft tissue of the brain deforms in response to external stimuli, which can lead to traumatic brain injury. Constitutive models relate the stress in the brain to its deformation and accurate constitutive modeling is critical in finite element simulations to estimate injury risk. Traditionally, researchers first choose a constitutive model and then fit the model parameters using tension, compression, or shear experiments. In contrast, constitutive artificial neural networks enable automated model discovery without having to choosing a specific model a priori before learning the model parameters. Here we reverse engineer a constitutive artificial neural network that uses the principal stretches, raised to a wide range of exponential powers, as activation functions for the hidden layer. Upon training, the network autonomously discovers a subclass of models with multiple Ogden terms that outperform popular constitutive models including neo Hooke, Blatz Ko, and Mooney Rivlin. While invariant-based networks fail to capture the pronounced tension-compression asymmetry of brain tissue, our principal-stretch-based network can simultaneously explain tension, compression, and shear data for the cortex, basal ganglia, corona radiata, and corpus callosum. Without fixing the number of terms a priori, our model self-selects the best subset of terms out of more than a million possible combinations, while simultaneously discovering the best model parameters and best experiment to train itself. Eliminating user-guided model selection has the potential to induce a paradigm shift in soft tissue modeling and democratize brain injury simulations.

Our source code, data, and examples are available at https://github.com/LivingMatterLab/CANN.

## Introduction

Understanding the mechanical behavior of brain tissue is essential to predict how the brain will respond when injured, during development, or how disease will progress [1]. Numerous constitutive models, which relate stress and stretch, have been developed from experimental data to best describe the behavior of soft tissues, including the brain [2]. These constitutive relations allow us to simulate how the brain will respond to various stress conditions, like traumatic brain injury, using techniques like the finite element method [3]. However, selecting the best constitutive model is not a trivial task as brain tissue is heterogeneous, highly nonlinear, displays pronounced tension-compression asymmetry, and is almost perfectly incompressible [4, 5].

### Simple loading experiments reveal the complex mechanical behavior of brain tissue

Simple loading experiments are frequently used to try to characterize the behavior of brain tissue [6]. However, because the brain tissue is ultrasoft, highly fragile, biphasic, heterogeneous, and deforms under its own weight due to gravity, getting accurate and reliable tissue responses from mechanical tests is difficult [6]. The most common mechanical tests include uniaxial tension [7], compression [8], shear [9], and indentation [10]. These tests have revealed that brain tissue has a strain-stiffening behavior and is stiffer under compression than tension [4]. Increasing the strain rate increases brain stiffness; the tissue also softens under preconditioning [6], but does not appear to be significantly anisotropic [4]. These results show just how complex the mechanical response of brain tissue is, and why it is so challenging to characterize its behavior with a single, simple constitutive equation.

### Constitutive artificial neural networks

Constitutive artificial neural networks show great promise in modeling the complex behaviors of soft biological tissues [11]. These networks have physics built into their architecture, and restricted inputs and outputs to follow the laws of physics and the governing principles for constitutive equations [12]. Critically, constitutive artificial neural networks do not require the user to *know* the form of the constitutive equation ahead of time [13]. Instead, they can *discover* the entire constitutive relationship automatically [12]. Previously, we have shown that constitutive artificial neural networks can be built from invariant-based building blocks like the neo Hooke, Blatz Ko, Demiray, and Holzapfel models [14]. While invariant-based neural networks perform well at fitting tension, compression, and shear individually, they struggle to capture the pronounced tension-compression asymmetry in brain tissue [14].

### Principal-stretch based models for soft tissue

The Ogden model for large deformations of incompressible rubber-like solids was first proposed more than half a century ago [15]. Its strain energy function features a linear combination of the principal stretches *λ*_1_, *λ*_2_, *λ*_3_ raised to arbitrary powers *α_k_* as 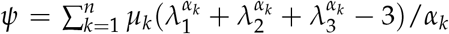. The Ogden model assumes that the material is isotropic [16], which makes it particularly well-suited for soft tissues such as the liver, kidney, blood clots, and the brain [17]. In most practical applications, the Ogden model is used with only one [9, 18, 19] or two terms [20] when fit to brain tissue data. A recent study showed that a subclass with up to eight Ogden terms provides excellent fits for both fat and brain tissue under combined compression and shear, while the common one- and two-term models provide inadequate fits [21]. However, identifiability becomes more difficult for these higher order Ogden models [5], and even the best algorithms struggle to fit these higher order models uniquely [22]. This raises the question to which extent higher order Ogden models are practically useful, and if so, whether and how we can identify them and fit them to data. The objective of this publication is to address this challenge by harnessing the power of gradient-based adaptive optimizers from deep learning and formulating the Ogden model as a constitutive artificial neural network that simultaneously learns the relevant Ogden terms and their parameters for human brain tissue.

## Methods

### Kinematics

During testing, particles ***X*** in the undeformed reference configuration map to particles ***x*** in the deformed configuration via the deformation map ***φ*** such that ***x*** = ***φ***(***X***). The deformation gradient ***F*** is the gradient of the deformation map ***φ*** with respect to the undeformed coordinates ***X***. Its spectral representation introduces the principal stretches *λ_i_* and the principal directions ***N**_i_* and ***n**_i_* in the undeformed and deformed configurations, where ***F*** · ***N**_i_* = *λ_i_**n**_i_*, and

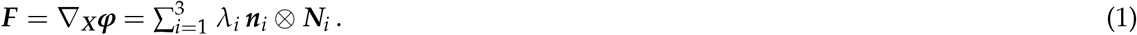

The Jacobian, *J* = det(***F***) > 0, denotes volumetric change, where 0 < *J* < 1 is compression, *J >* 1 is expansion, and *J* = 1 means the material is *perfectly incompressible*. To measure the deformation in the undeformed configuration, we introduce the right Cauchy Green tensor **C** and its spectral representation in terms of principal stretches squared 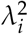 and the principal directions ***N**_i_*,

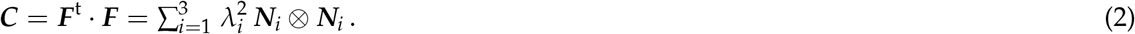

An *isotropic* material has three principal invariants,

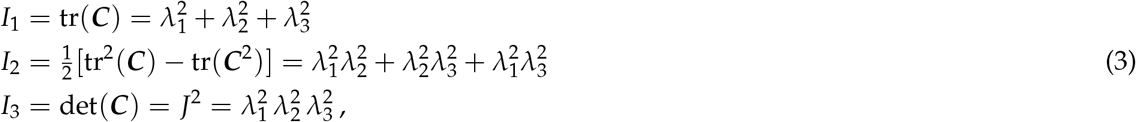

which are linear, quadratic, and cubic in terms of these principal stretches squared. The principal stretches depend on the type of experiment. For *uniaxial tension and compression* tests, where we apply a stretch *λ*, we can write the deformation gradient ***F*** in matrix representation as

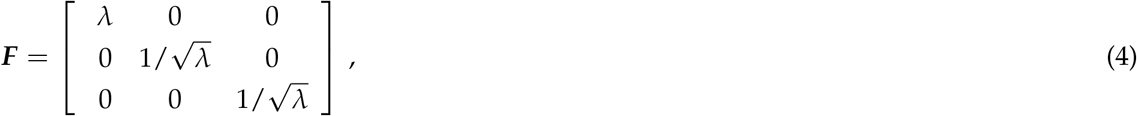

and calculate the principal stretches as

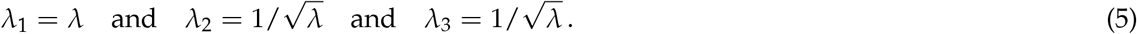

For *simple shear* tests where we apply a shear *γ*, the deformation gradient **F** in matrix representation is

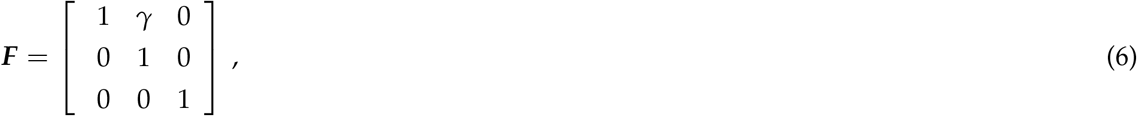

and the principal stretches are

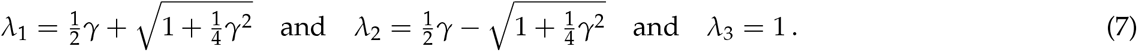

### Constitutive equations

Constitutive equations relate a stress like the Piola or nominal stress ***P***, the force per undeformed area that is commonly measured in experiments, to a deformation measure like the deformation gradient ***F***. For a *hyperelastic* material that satisfies the second law of thermodynamics, we can express the Piola stress, ***P*** = *∂ψ*(***F***)/*∂**F***, as the derivative of the Helmholtz free energy function *ψ*(***F***) with respect to the deformation gradient ***F***, modified by a pressure term, –*p **F***^-t^, to ensure *perfect incompressibility*,

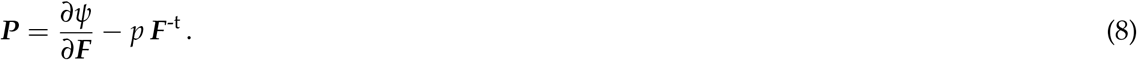

Instead of formulating the free energy function in terms of the deformation gradient, several constitutive models define the free energy function in terms of the three principal stretches *λ_i_*. For an *isotropic* material, we can express the Piola stress, 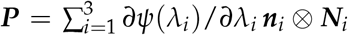, as the derivative of the Helmholtz free energy function *ψ*(*λ_i_*) with respect to the principal stretches *λ_i_*, modified by the pressure term,

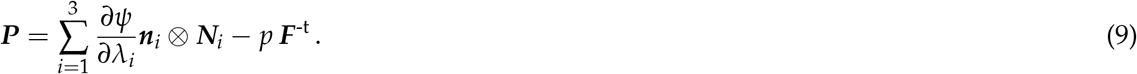

The hydrostatic pressure, 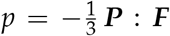 acts as a Lagrange multiplier that that we determine from the boundary conditions.

### Principal-stretch-based model

The free energy function of the Ogden model [15] is a function of the three principal stretches, *λ*_1_, *λ*_2_, *λ*_3_,

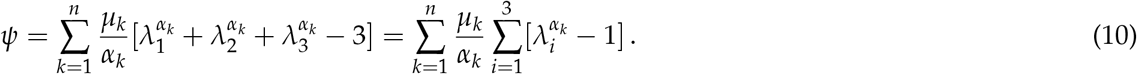

It consists of *n* terms and introduces 2*n* parameters, *n* stiffness-like parameters *μ_i_* and *n* nonlinearity parameters *α*_i_, which collectively translate into the classical shear modulus *μ* from linear theory [2],

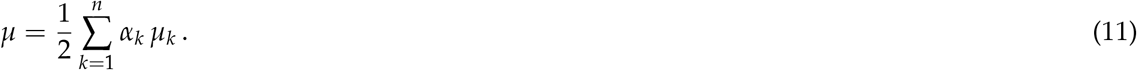

### Principal-stretch-based constitutive artificial neural network

Based on these guiding physical principles, we reverse-engineer a hyperelastic, perfectly incompressible, isotropic, principal-stretch-based constitutive artificial neural network. The network takes the deformation gradient ***F*** as input and then computes the principal stretches *λ*_1_, *λ*_2_, *λ*_3_. From these stretches, it determines *n* Ogden terms, with fixed exponential coefficients, here ranging from *α*_1_ = −30 to *α_n_* = +10 in increments of two, that make up the *n* nodes of the hidden layer of the model. The weighted sum of all terms, defines the strain energy function *ψ* as the network output.

Figure 1 illustrates a principal-stretch-based constitutive artificial neural network with *n* = 20 nodes, for which the free energy function takes the following form,

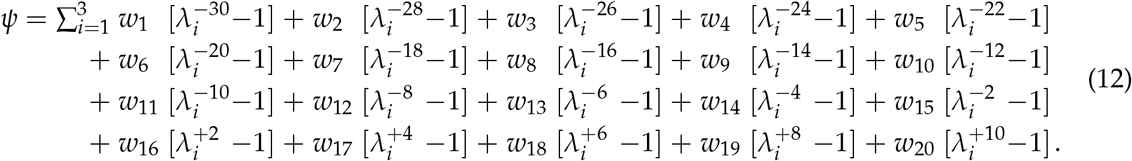

**Figure 1:**
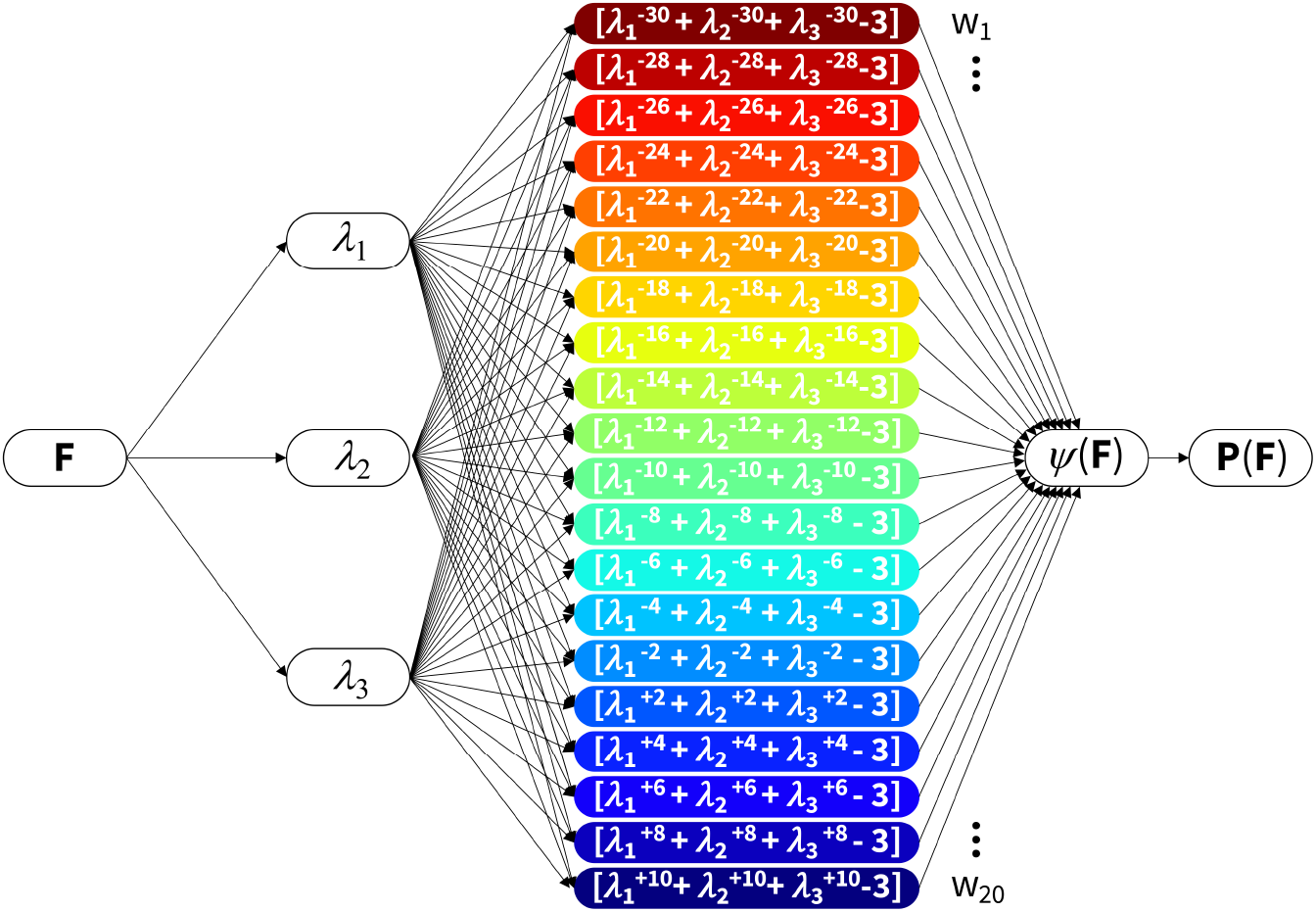
Principal-stretch-based constitutive artificial neural network. The network represents a 20-term Ogden model with fixed exponents *α_k_* ranging from −30 to +10 in increments of two. It takes the deformation gradient ***F*** as input and computes the three principal stretches *λ*_1_, *λ*_2_, *λ*_3_. From them, it calculates the Ogden terms for the 20 nodes of the hidden layer, multiplies each nodal term by the network weight *w_k_*, and adds all terms to the strain energy function *ψ*(***F***) as output. The derivative of the strain energy function defines the Piola stress, **P** = *∂ψ/∂***F**, whose components *P*_11_ or *P*_12_ enter the loss function to minimize the error with respect to the tension, compression, and shear data.

Here, *w_k_* are the network weights and 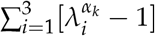 are the activation functions. Figure 2 illustrates the *n* = 20 activation functions of our neural network for the load cases of uniaxial tension and compression in the top block and simple shear in the bottom block. From the free energy (12), we calculate the stress using equation (9),

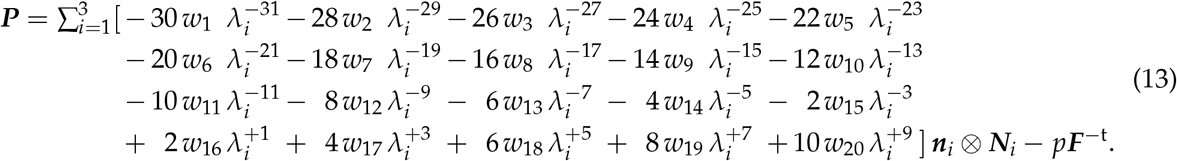

**Figure 2:**
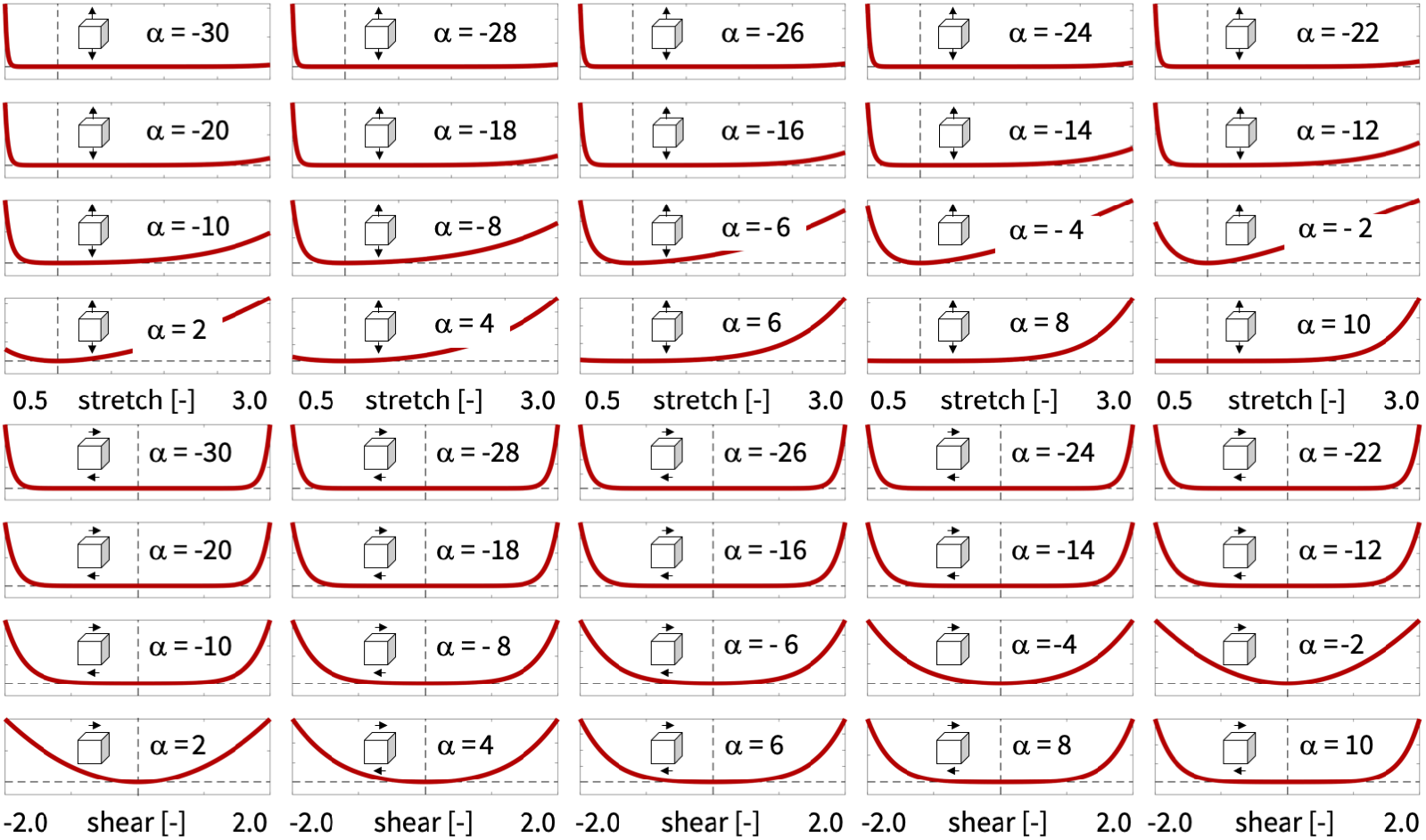
Activation functions of principal-stretch-based constitutive artificial neural network. The activation functions 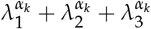 represent the contributions to the free energy function *ψ* of a 20-term Ogden model with fixed exponents *α_k_* ranging from −30 to +10 in increments of two. Activation functions in the top block are associated with uniaxial tension and compression, with principal stretches 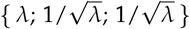 plotted over the range 0.5 ≤ *λ* ≤ 3.0; activation functions in the bottom block are associated with simple shear, with principal stretches 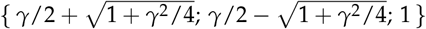 plotted overthe range −2.0 ≤ *γ* ≤ 2.0. The sum of all activation functions, weighted by the network weights *w_k_*, represents the free energy function *ψ* of the Ogden model.

The network weights *w_k_* are non-negative [13], and relate to the stiffness-like parameters *μ_k_* and fixed exponential coefficients *α_k_* as

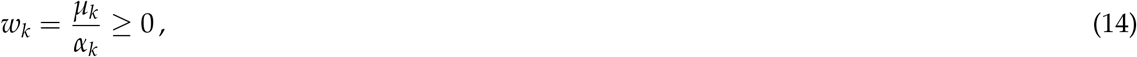

where, for our specific network, the coefficients *α_k_* are *α_k_* = 2*k* – *n* – 12 for *k* ≤ *n* – 5 and *α_k_* = 2*k* – *n* – 10 for *k* ≥ *n* – 5. For this particular network, we can express the classical shear modulus *μ* from the linear theory (11) in terms of the stiffness-like parameters *μ_k_*,

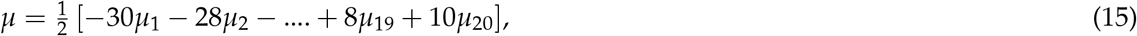

or, equivalently, in terms of the network weights *w_k_*,

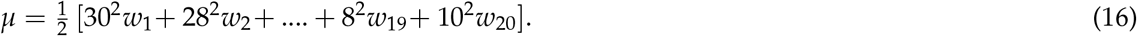

Notably, unlike conventional neural networks that use *similar* activation functions at all nodes, our constitutive artificial neural network uses *different* activation functions at all *n* nodes, as indicated in Figure 2. During training, our network autonomously identifies the best subset of activation functions from (2*^n^* – 1) = 1, 048, 575 possible combinations, and discovers the best model from more than a million possible models.

### Special cases

Our principal-stretch-based constitutive artificial neural network in Figure 1 is a generalization of popular constitutive models. Specifically, we obtain the following one- and two-term Ogden models by setting the remaining 19 and 20 network weights to zero.

The *neo Hooke* model [23] only uses the fixed exponent *α*_16_ = +2 and its free energy function is

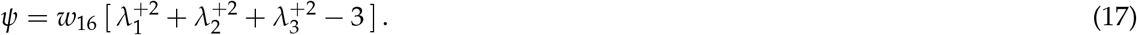

Here, *w*_16_ = *μ*_16_/2 and *μ* = *μ*_16_. So, the shear modulus of the neo Hooke model is *μ* = 2 *w*_16_.

For the *Blatz Ko* model [24] only uses the fixed exponent *α*_15_ = −2 and its free energy function is

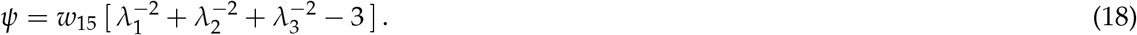

Here, *w*_15_ = −*μ*_15_/2 and *μ* = −*μ*_15_. So, the shear modulus of the Blatz Ko model is *μ* = 2*w*_15_.

The *Mooney Rivlin* model [25, 26] only uses the fixed exponents *α*_15_ = −2 and *α*_16_ = +2 and its free energy function is

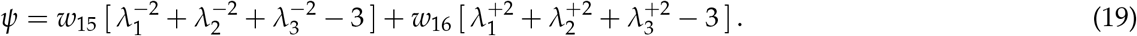

Here, w_15_ = –*μ*_15_/2, w_16_ = *μ*_16_/2, and *μ* = – *μ*_15_ + *μ*_16_. So, the shear modulus of the Mooney Rivlin model is *μ* = 2 [*w*_15_ + *w*_16_].

The *general one-term Ogden model* [15] only uses a single free exponents *α* and its free energy function is

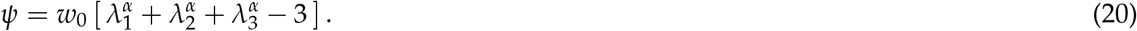

Here, *w*_0_ = *μ*_0_/*α* and *μ* = *αμ*_0_/2. So the shear modulus of the general one-term Ogden model is *μ* = *α*^2^w_0_/2.

### Loss function

Our constitutive artificial neural network learns the network weights, ***w*** = *w*_1_,…, *w_k_*, by minimizing a loss function *L* that penalizes the mean squared error, the *L*_2_-norm of the difference between model ***P***(***F**_i_*) and data 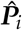, divided by the number of training points *n*_train_,

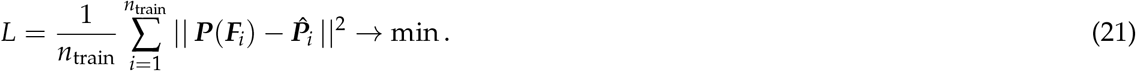

To reduce potential overfitting, we also study the effects of Lasso or L1 regularization,

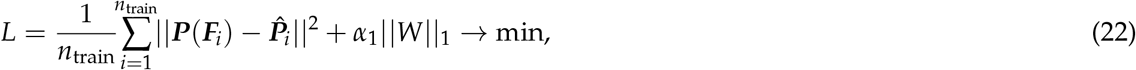

where *α*_1_ is the penalty parameter or regularization coefficient and ∥*W*∥_1_ = ∑*_k_*|*w_k_*| is the weighted L1 norm. We train the network by minimizing the loss functions (21) or (22) and learn the network parameters *w_k_* using the ADAM optimizer, a robust adaptive algorithm for gradient-based first-order optimization, and constrain the weights to always remain non-negative, *w_k_* ≥ 0.

### Data

We train and test our principal-stretch-based constitutive artificial neural network using tension, compression, and shear data of human gray matter tissue from the cortex and basal ganglia and white matter tissue from the corona radiata and corpus callosum [4], as reported in Table 1. [14]. We perform *single-mode training* using a single loading case, either tension, compression, or shear, as training data and the remaining two cases as test data. We also perform *multi-mode training* using all three loading cases simultaneously as training data.

**Table 1:**
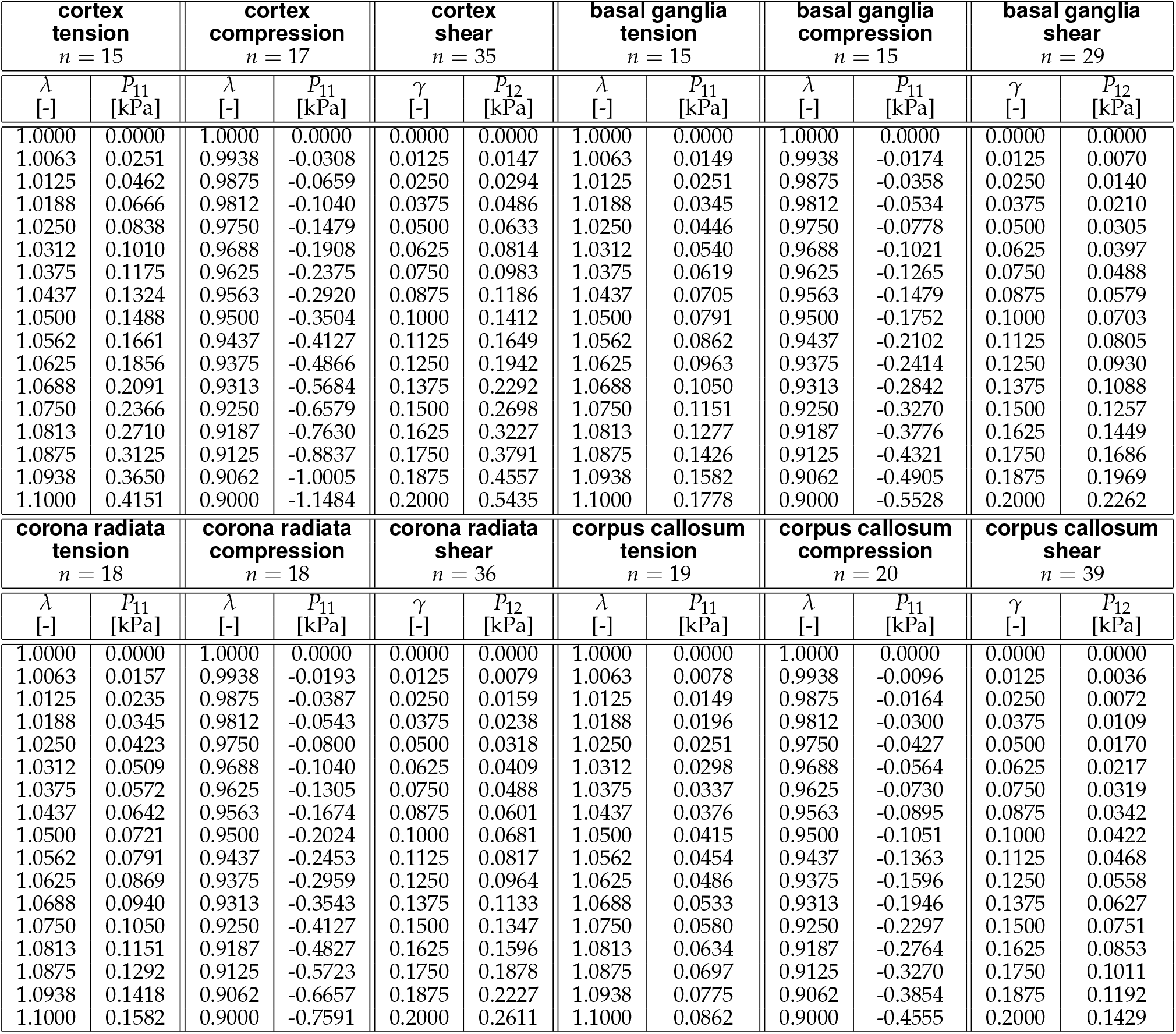
Cortex, basal ganglia, corona radiata, and corpus callosum tested in tension, compression, and shear. Stresses are reported as means from the loading and unloading curves of *n* samples tested in the ranges 1.0 ≤ *λ* ≤ 1.1 for tension, 0.9 ≤ *λ* ≤ 1.0 for compression, and 0.0 ≤ *γ* ≤ 0.2 for shear [4].

## Results

Figure 3 illustrates the automatically discovered constitutive models for the human cortex using the twenty-term isotropic, perfectly incompressible, principal-stretch-based constitutive artificial neural network. It shows the Piola stress as a function of the stretch or shear strain. The first three columns indicate single-mode training for tension, compression, or shear, training on the diagonal and testing on the off-diagonal. The last column shows the results of multi-mode training with all three loading modes as training data. The circles indicate the experimental data from Table 1, while the color-coded regions designate the contributions of the twenty model terms to the free energy function *ψ*. Warm red-type colors indicate negative exponents, while cold blue-type colors indicate positive exponents. Each graph reports the coefficient of determination, *R*^2^, as a measure for the goodness of fit. First, for single-mode training, our principal-stretch-based constitutive artificial neural network succeeds in *fitting* the individual sets of training data with 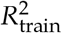 values of 0.9085, 0.9997, and 0.9994 for tension, compression, and shear. Second, for single-mode training, the network performs moderately well in *predicting* the test data with 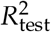 values ranging from 0.1753 for the compression prediction with tension training to 0.9735 for the shear prediction with compression training. Third, for multi-mode training, the training fit 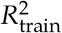 slightly decreases for compression to 0.9851 but increases for tension and shear to 0.9377 and 0.9873. Overall, the sum of all three *R*^2^ values increases significantly compared to single-mode training. Fourth, for single-mode training, the network finds a wide spectrum of non-zero terms ranging from dark red with *α* = −30 to dark blue with *α* = +10, while for multimode training, the red-type negative exponential terms dominate over the blue-type terms.

**Figure 3:**
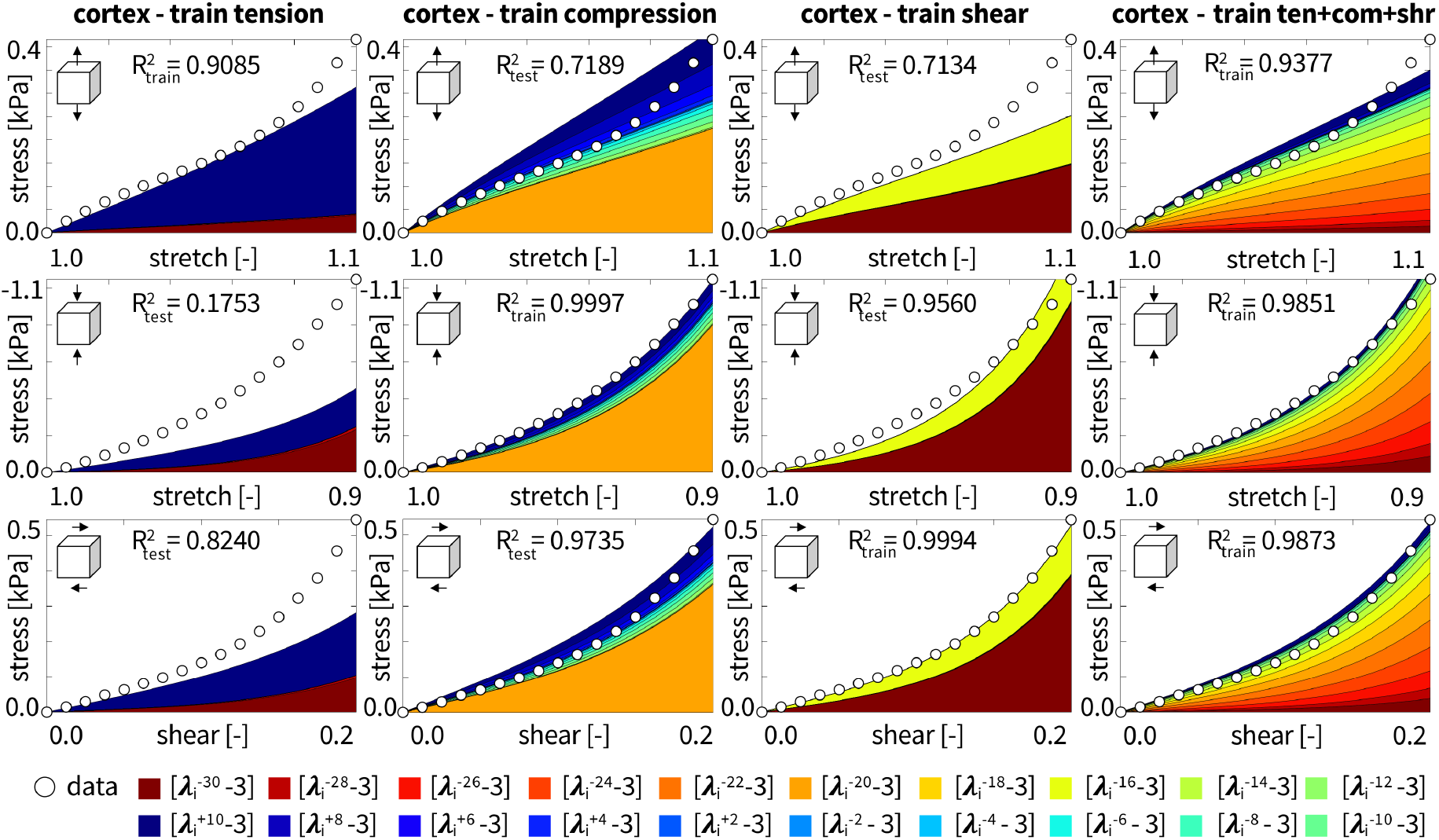
Cortex data and model. Nominal stress as a function of stretch or shear strain for the principal-stretch-based constitutive artificial neural network with one hidden layer and twenty nodes from Figure 1. Training individually with tension, compression, shear data from the human cortex, and with all three load cases simultaneously. Circles represent the experimental data. Color-coded regions designate the contributions of the 20 model terms to the stress function according to Figure 1. Coefficients of determination R^2^ indicate goodness of fit for train and test data.

Figure 4 illustrates the automatically discovered models for the human corona radiata for the tension, compression, and shear data. The corona radiata is part of the white matter in the brain. Its results confirm similar models trends to that of the grey matter results for the cortex in Figure 3. First, for single-mode training, the network succeeds in *fitting* the individual sets of training data with 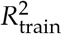 values of 0.9626, 0.9957, and 0.9994 for tension, compression, and shear. Second, for single-mode training, the network performs moderately well in *predicting* the test data with 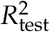 values ranging from 0.4251 for the tension prediction with compression training to 0.9176 for the shear prediction with compression training. Third, for multi-mode training, the training fit 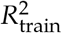 slightly decreases for all three loading modes to 0.9453, 0.9889, and 0.9388 for tension, compression, and shear. However, the sum of all three *R*^2^ values increases significantly compared to the single-mode training. Fourth, for single-mode training, the network finds a wide spectrum of terms ranging from dark red with *α* = −30 to dark blue with *α* = +10, while for multi-mode training, it primarily discovers the red-type negative exponential terms.

**Figure 4:**
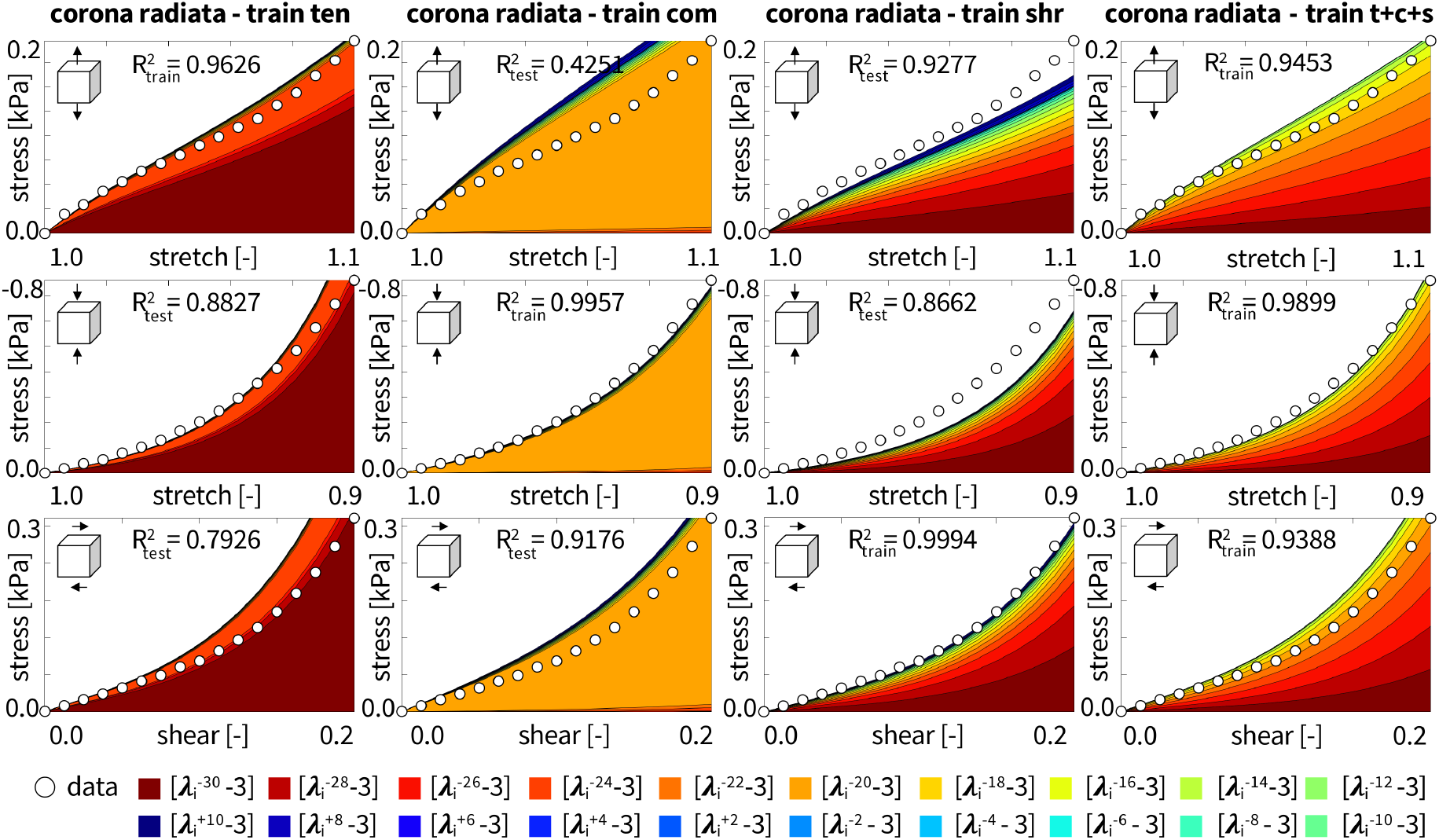
Corona radiata data and model. Nominal stress as a function of stretch or shear strain for the principal-stretchbased constitutive artificial neural network with one hidden layer and twenty nodes from Figure 1. Training individually with tension, compression, shear data from the human corona radiata, and with all three load cases simultaneously. Circles represent the experimental data. Color-coded regions designate the contributions of the 20 model terms to the stress function according to Figure 1. Coefficients of determination R^2^ indicate goodness of fit for train and test data.

Figure 5 and Table 2 summarize the automatically discovered models for the human cortex, basal ganglia, corona radiata, and corpus callosum in multi-mode training on tension, compression, and shear data. First, the fit for the compression data is the best across all four brain regions with 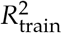 values of 0.9851, 0.9803, 0.9899, and 0.9592. However, the fits for the tension and shear are also comparable, with 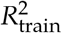 values ranging from 0.9114 for the corpus callosum in tension to 0.9946 for the basal ganglia in shear. Second, the red-type color terms with negative exponents dominate over the positive blue-type terms for all brain regions. This confirms the trends in Figures 3 and 4. Third, from the discovered weights in Table 2, we can derive the overall shear moduli of 1.4753 kPa, 0.6828 kPa, 0.6940 kPa, and 0.2933 kPa for the cortex, basal ganglia, corona radiata, and corpus callosum suggesting that the cortex is the stiffest region, followed by both the basal ganglia and the corona radiata, which are approximately half as stiff as the cortex, and the corpus callosum, which is in turn half as stiff as these two regions. These findings agree well with the shear moduli of 1.43 kPa, 0.70 kPa, 0.66 kPa, and 0.35 kPa fitted for a one-term Ogden model with all three loading modes [4]. Notably, this study identified single-term exponential powers *α* of −43.6, −32.5, −30.5, and −26.6 for the four regions. Trained on the same data, our invariant-based neural network discovered moduli of 1.82 kPa, 0.88 kPa, 0.94 kPa, and 0.54 kPa for these four brain regions [14].

**Figure 5:**
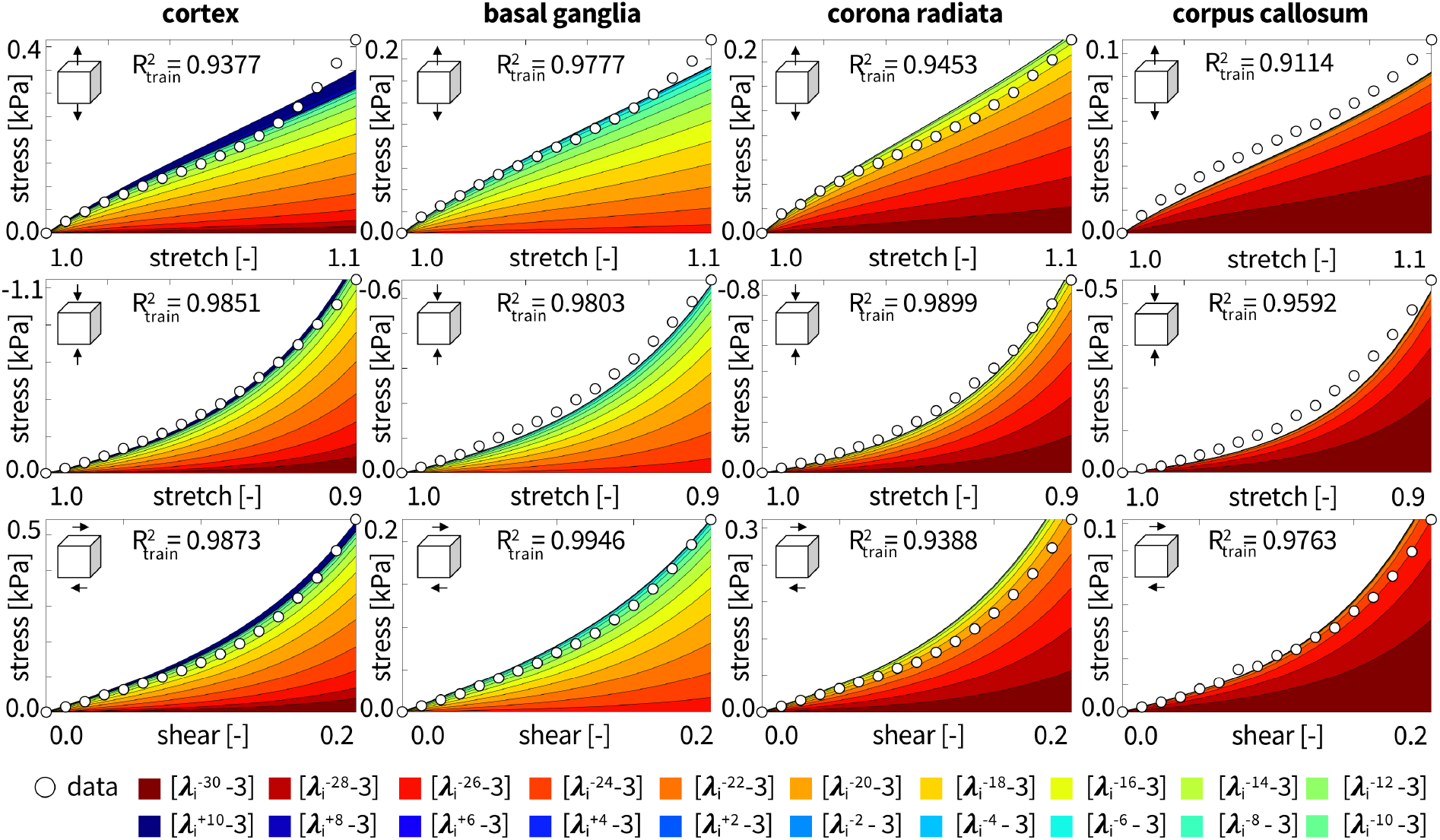
Cortex, basal ganglia, corona radiata, and corpus callosum data and models. Nominal stress as a function of stretch or shear strain for the principal-stretch-based constitutive artificial neural network with one hidden layer and twenty nodes from Figure 1. Training with tension, compression, shear data simultaneously. Circles represent the experimental data. Color-coded regions designate the contributions of the 20 model terms to the stress function according to Figure 1 multiplied by the weights from Table 2. Coefficients of determination R^2^ indicate goodness of fit for train data.

**Table 2:** Cortex, basal ganglia, corona radiata, and corpus callosum models and parameters. Models and parameters are discovered for simultaneous training with tension, compression, and shear data using the principal-stretch-based constitutive artificial neural network with twenty nodes from Figure 1. Summary of the weights *w_k_*, shear moduli *μ* calculated from equation (16), and goodness of fit *R*^2^ for training in tension, compression, and shear.

| | **cortex**<br>**ten+com+shr**<br>$n = 15, 17, 35$ | **basal ganglia**<br>**ten+com+shr**<br>$n = 15, 15, 29$ | **corona radiata**<br>**ten+com+shr**<br>$n = 18, 18, 36$ | **corpus callosum**<br>**ten+com+shr**<br>$n = 19, 20, 39$ |
| --- | --- | --- | --- | --- |
|  | [kPa] | [kPa] | [kPa] | [kPa] |
| $w_1$ | 0.000127 | 0.000000 | 0.000189 | 0.000227 |
| $w_2$ | 0.000128 | 0.000000 | 0.000252 | 0.000199 |
| $w_3$ | 0.000292 | 0.000097 | 0.000307 | 0.000164 |
| $w_4$ | 0.000507 | 0.000229 | 0.000350 | 0.000120 |
| $w_5$ | 0.000781 | 0.000355 | 0.000378 | 0.000068 |
| $w_6$ | 0.000991 | 0.000472 | 0.000386 | 0.000015 |
| $w_7$ | 0.001132 | 0.000576 | 0.000370 | 0.000010 |
| $w_8$ | 0.001200 | 0.000670 | 0.000326 | 0.000005 |
| $w_9$ | 0.001196 | 0.000748 | 0.000248 | 0.000005 |
| $w_{10}$ | 0.001120 | 0.000802 | 0.000132 | 0.000005 |
| $w_{11}$ | 0.000978 | 0.000827 | 0.000018 | 0.000005 |
| $w_{12}$ | 0.000779 | 0.000852 | 0.000012 | 0.000004 |
| $w_{13}$ | 0.000583 | 0.000847 | 0.000006 | 0.000003 |
| $w_{14}$ | 0.000382 | 0.000800 | 0.000001 | 0.000002 |
| $w_{15}$ | 0.000239 | 0.000711 | 0.000000 | 0.000000 |
| $w_{16}$ | 0.000235 | 0.000399 | 0.000000 | 0.000000 |
| $w_{17}$ | 0.000244 | 0.000179 | 0.000000 | 0.000000 |
| $w_{18}$ | 0.000265 | 0.000004 | 0.000000 | 0.000000 |
| $w_{19}$ | 0.000826 | 0.000000 | 0.000000 | 0.000000 |
| $w_{20}$ | 0.001594 | 0.000000 | 0.000000 | 0.000000 |
| | $\mu = 1.4753$ kPa | $\mu = 0.6828$ kPa | $\mu = 0.6940$ kPa | $\mu = 0.2933$ kPa |
| $R_t^2$ | 0.9377 | 0.9777 | 0.9453 | 0.9114 |
| $R_c^2$ | 0.9851 | 0.9803 | 0.9899 | 0.9592 |
| $R_s^2$ | 0.9873 | 0.9946 | 0.9388 | 0.9763 |

Figure 6 illustrates the special case of the one-term principal-stretch-based model related to the −24, −22, −20, and −18 terms, in multi-mode training for the human cortex. These four terms are the terms with the largest contributions to the stress in Figure 3. Visually, this means they correspond to the dominant colors in the graphs. First, the −18 term provided the best overall fits with 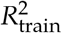 values of 0.9123, 0.9858, and 0.9871. Second, the best single-term model with −18 was not as good at fitting the data as the discovered model in Figure 3 with the biggest decrease in performance for the tension fit. Third, from Table 3 the shear moduli were 1.0294 kPa, 1.3130 kPa, 1.3263 kPa, and 1.4010 kPa for the −24, −22, −20, and −18 term models. The modulus for the −18 term is comparable to the reported modulus of 1.43 kPa identified for the one-term Ogden model trained on all three loading modes with *α* = −19.0 [4]. These results suggest that the one-term principal-stretch-based models tend to underestimate the shear modulus of brain tissue compared to multi-term models.

**Figure 6:**
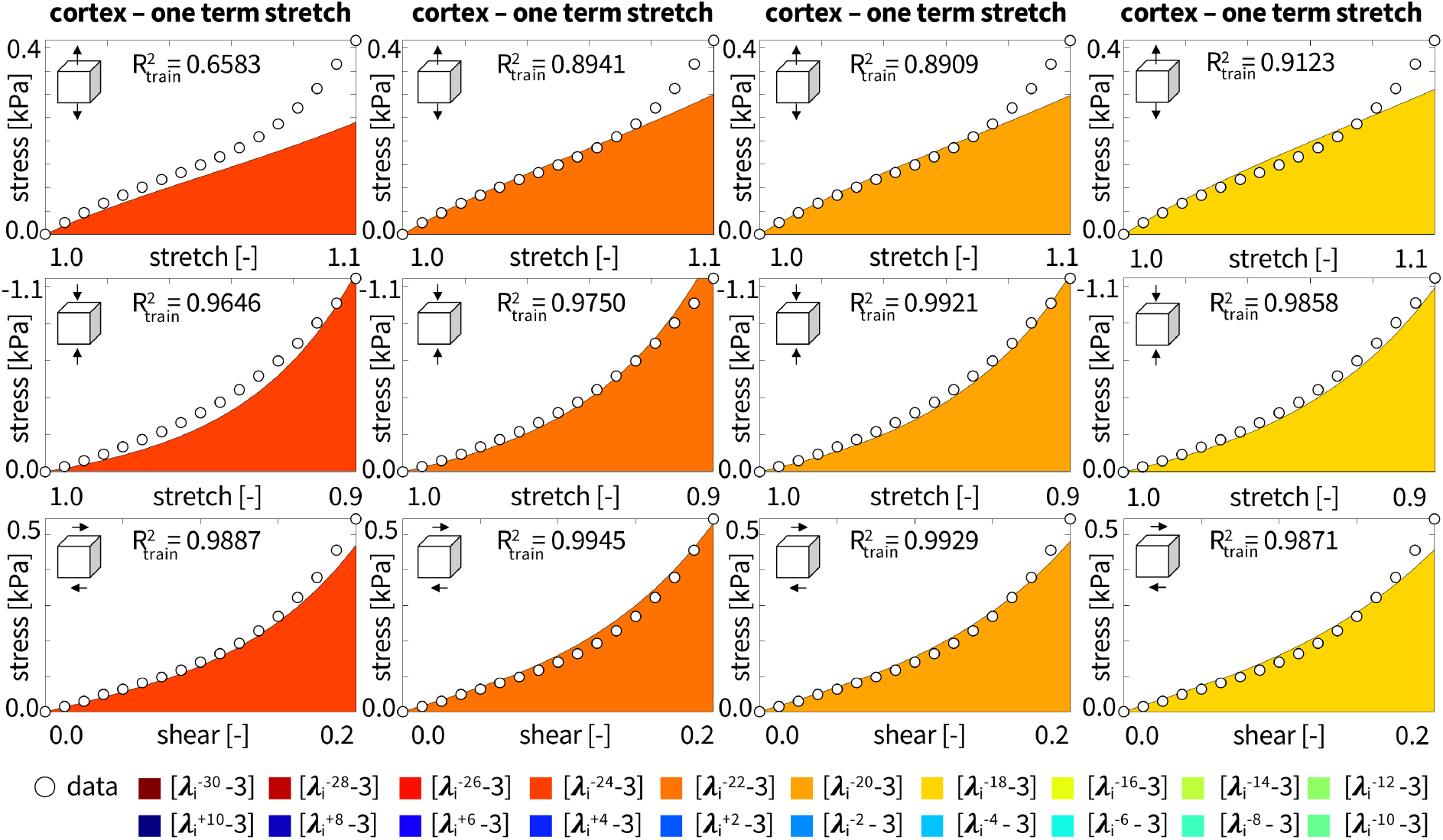
Cortex data and special case of one-term models with −24, −22, −20, −18 terms. Nominal stress as a function of stretch or shear strain for the principal-stretch-based one-term model with −24, −22, −20, −18 terms, selected as the terms with the largest pressure contribution in Figure 3 when trained with all three loading cases from the human cortex simultaneously. Circles represent the experimental data. Color-coded regions designate the −24, −22, −20, −18 model terms to the stress function according to Figure 1. Coefficients of determination R^2^ indicate goodness of fit for train data.

**Table 3:**
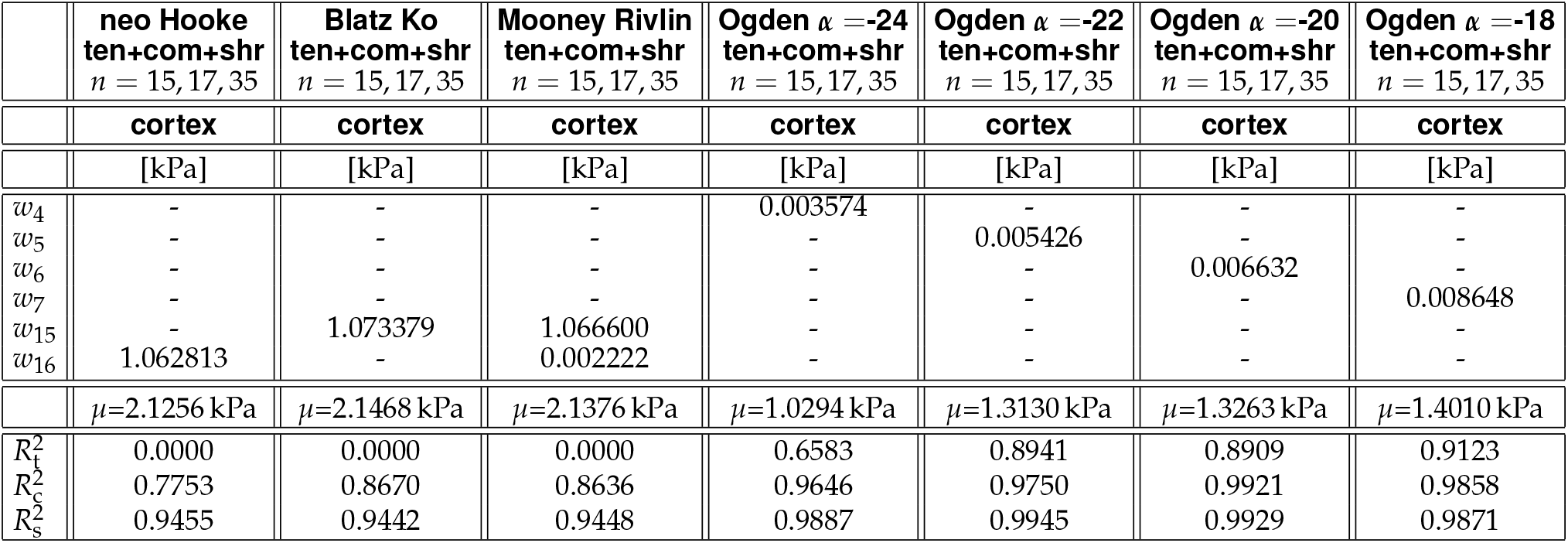
Cortex special cases of neo Hooke, Blatz Ko, Mooney Rivlin, and one-term Ogden models and parameters. Cortex parameters discovered for simultaneous training with tension, compression, and shear data. Summary of the nonzero weights, shear moduli *μ*, and goodness of fit *R*^2^ for the training data.

Figure 7 illustrates four special cases, the neo Hooke, Blatz Ko, Mooney Rivlin, and one-term Ogden models, in multi-mode training for the human cortex. The neo Hooke model uses the +2 term, the Blatz Ko the −2 term, the Mooney Rivlin both the +2 and −2 terms, and the one-term Ogden model the −18 term from Figure 6. First, the commonly used neo Hooke, Blatz Ko, and Mooney Rivlin models all fail to explain the tension data with 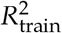 values of 0.0000. Second, the neo Hooke, Blatz Ko, and Mooney Rivlin models are able to provide adequate fits for the cortex data in compression and shear with 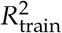 values ranging from 0.7753 to 0.9444. Third, the one-term Ogden model from the fourth column of Figure 6 outperforms the neo Hooke, Blatz Ko, and Mooney Rivlin models for all three loading modes. Fourth, the shear moduli calculated from the weights was 2.1256 kPa, 2.1468 kPa, 2.1376 kPa, and 1.4010 kPa for the neo Hooke, Blatz Ko, Mooney Rivlin, and one-term Ogden model with the −18 term. The neo Hooke and Mooney Rivlin shear moduli agree well with the values of 2.07 kPa and 2.08 kPa reported for the cortex with simultaneous fitting on all three loading modes [4].

**Figure 7:**
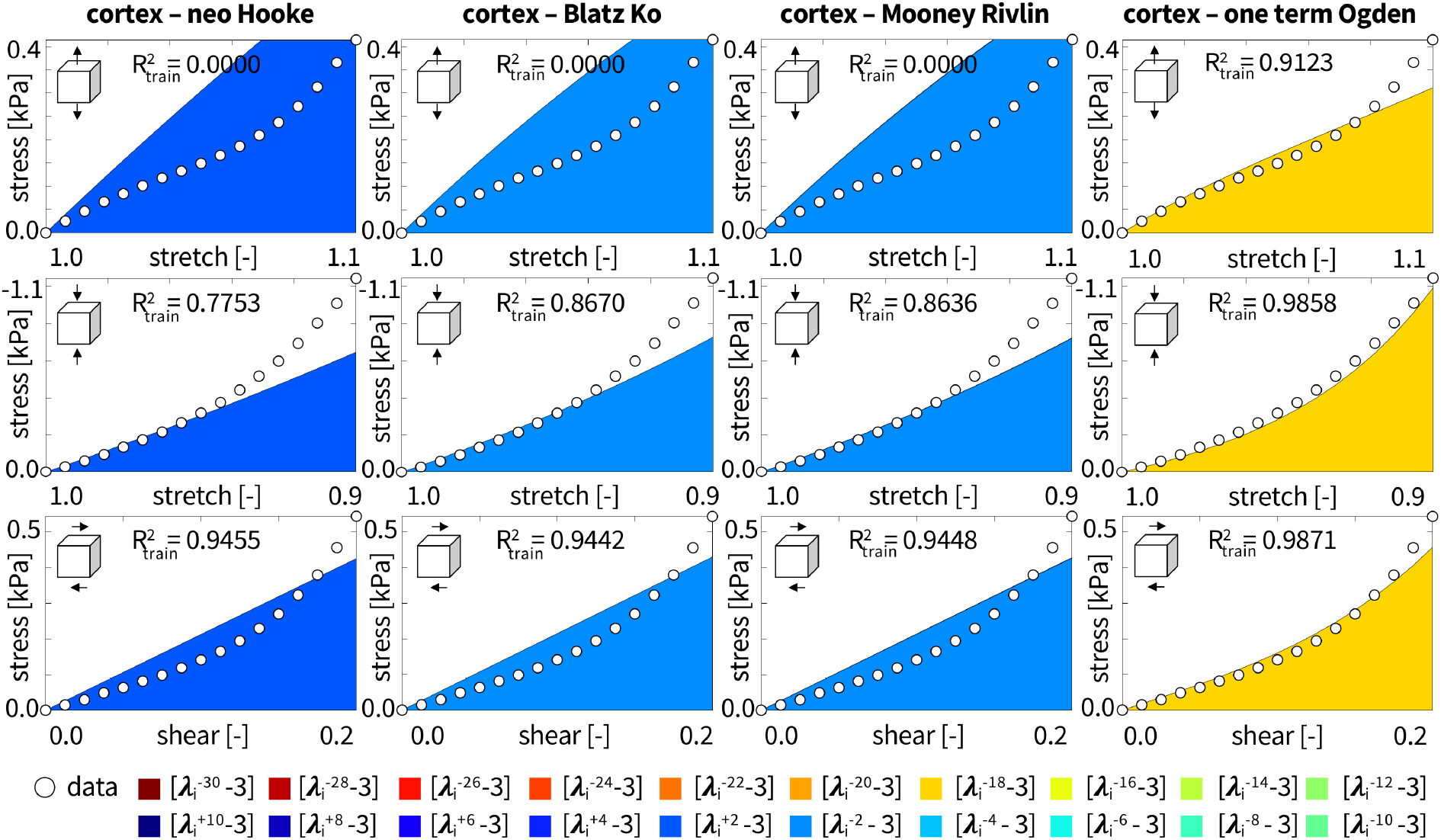
Cortex data and special cases of neo Hooke, Blatz Ko, Mooney Rivlin, and one-term Ogden models. Nominal stress as a function of stretch or shear strain for the principal-stretch-based models when trained with all three loading cases from the human cortex simultaneously. The neo Hooke model uses only the +2 term, the Blatz Ko the −2 term, the Mooney Rivlin the −2 and +2 terms, and the one-term Ogden the −18 term. Color-coded regions designate the +2, −2, +2/-2, −18 model terms to the stress function according to Figure 1 multiplied by the weights from Table 3. Coefficients of determination R^2^ indicate goodness of fit for train data.

Figure 8 illustrates the effect of added L1 regularization in multi-mode training for the human cortex. The L1 penalty parameters are 10, 1, 0.1, and 0. First, the moderately penalized models with 1 and 0.1 provide a good fit for all three loading cases, with 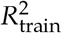 ranging from 0.8774 for the penalty parameter of 1 in tension to 0.9917 for the penalty parameter of 0.1 in shear. Second, similar to the non-regularized multi-mode training in Figures 3, 4, and 5, the red-type negative exponents dominate over the positive blue-type terms, especially with an increasing penalty parameter. Third, the largest L1 penalty parameter of 10 decreases the number of discovered terms the most, to only one, suggesting this model is over-regularized. The penalty parameter of 0.1 discovers 15 terms, five terms less terms than the non-regularized model, indicating that moderate L1 regularization successfully decreases the number of discovered terms.

**Figure 8:**
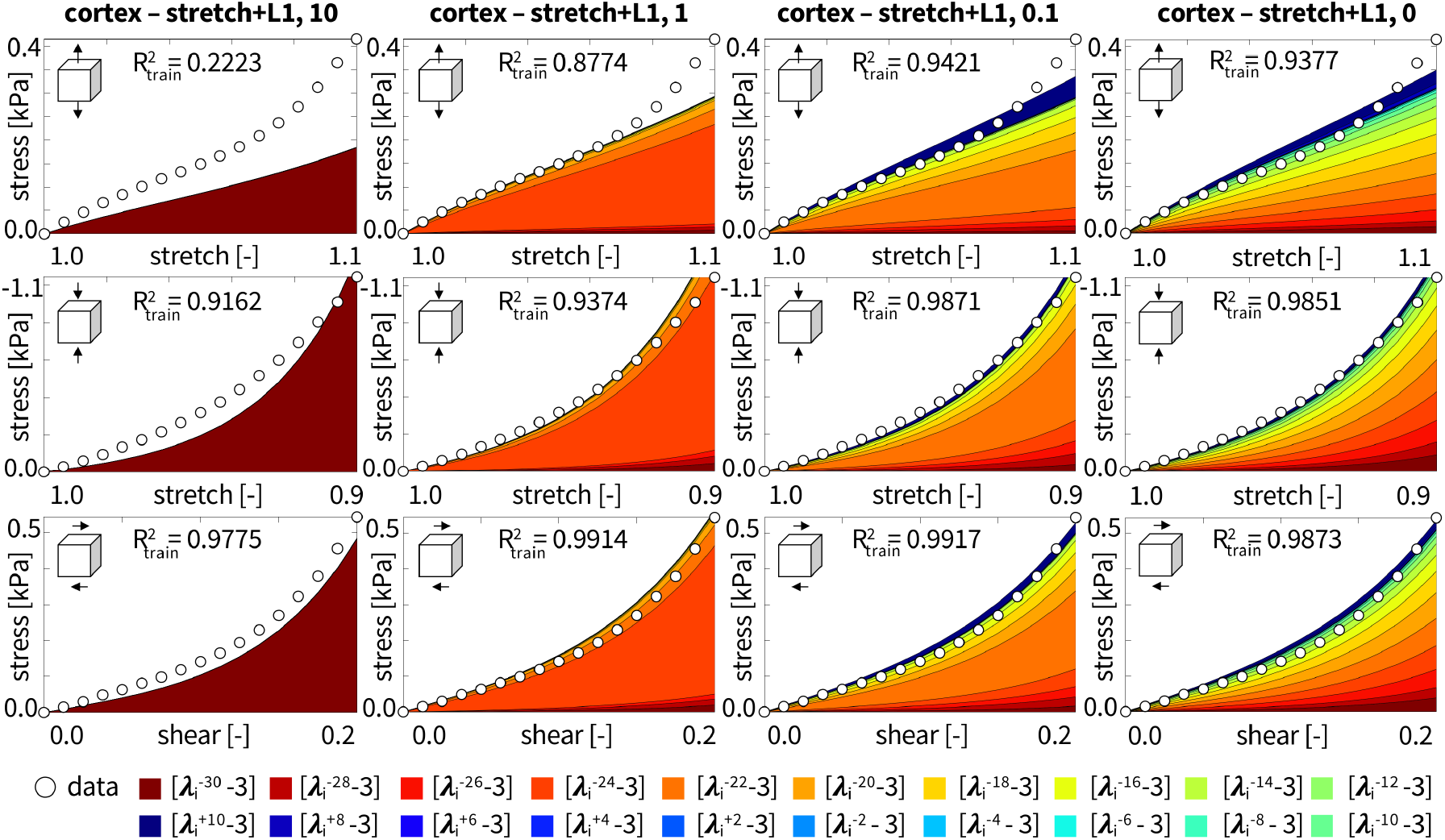
Effect of L1 regularization for varying penalty parameters. Nominal stress as a function of stretch or shear strain for the principal-stretch-based constitutive artificial neural network with one hidden layer and twenty nodes from Figure 1. L1 regularization is added to the loss function with regularization coefficients of 10, 1, 0.1, and 0. Training with tension, compression, shear data from the human cortex simultaneously. Circles represent the experimental data. Color-coded regions designate the contributions of the 20 model terms to the stress function according to Figure 1. Coefficients of determination R^2^ indicate goodness of fit for train data.

Figure 9 compares the performance of the neo Hooke, Blatz Ko, Mooney Rivlin, and principal-stretch-based constitutive artificial neural network without and with L1 regularization. For comparison, we also include the invariant-based constitutive artificial neural network without and with L2 regularization [14]. The bars represent the means and standard deviations of the coefficients of determination *R*^2^ across all four brain regions. The first three columns are the results of single-mode training on tension, compression, and shear with training on the diagonal and testing on the off-diagonal. The last column is the result of multi-mode training on all three loading modes simultaneously. First, both principal-stretch-based networks outperform both invariant-based networks for tension in multi-mode training and in all cases except for tension and shear predictions with tension training where the error bars are still within the same range. For multi-mode training in tension, the *R*^2^ values more than double. Notably, the principal-stretch-based models with and without L1 regularization are the *only* models capable of predicting tension from compression training. Second, *regularization improves generalization* regardless of whether the network is invariant-based or principal-stretch-based. The L2 penalty parameter was 0.001 for the invariant-based model, and the L1 penalty parameter was 1 for the principal-stretch-based model. Both L1 and L2 regularized networks produced better fits than their non-regularized counterparts for shear testing with compression training. Notably, the L1-regularized principal-stretch-based network also performed better than its non-regularized counterpart for the tension data with compression training. Third, the neo Hooke, Blatz Ko, and Mooney Rivlin models showed similar results to each other and to both invariant-based networks. Compared to both principal-stretch-based networks, however, these special cases did not perform as well for the tension fit with multi-mode training, compression fit with tension or shear training, and tension fit with compression training. Finally, we conclude that both principal-stretch-based networks provide the best fit across all three loading modes and all four brain regions with multi-mode training. Similarly to the invariant-based model, we notice that the principal-stretch-based networks consistently show high R^2^ values and low standard deviations across the shear row, indicating that *shear is the best experiment* to fit and predict with. For single-mode training, the most informative test for the non-regularized principal-stretch-based network is the shear test with the largest R^2^ values across all three loading modes. Interestingly, for the L1 regularized principal-stretch-based network, compression is the most informative test. Taken together, the *principal-stretch-based network outperforms the invariant-based network* in seven cases, does equally as well in two cases, and does slightly worse in three. For single- or multi-mode training, the principal-stretch-based constitutive artificial neural network is clearly the best choice of model.

**Figure 9:**
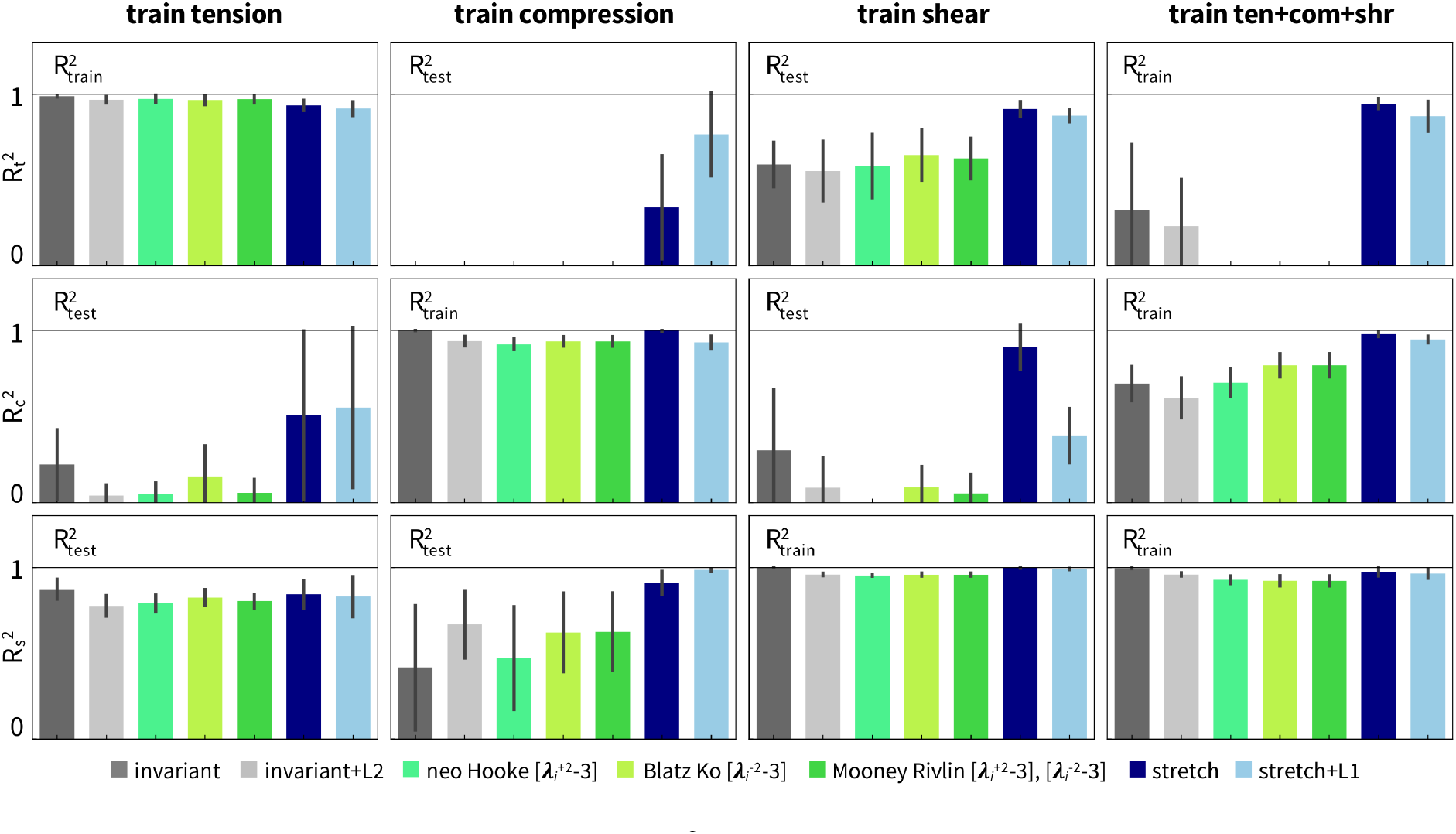
Goodness of fit for all seven models. Mean R^2^ and standard deviation for the four brain regions for each model, invariant-based constitutive artificial neural network without and with L2 regularization, neo Hooke, Blatz Ko, Mooney Rivlin, and principal-stretch-based constitutive artificial neural network without and with L1 regularization. The neo Hooke, Blatz Ko, and Mooney Rivlin models represent special cases of the principal-stretch-based constitutive artificial neural network with +2, −2, +2/-2. Rows correspond to tension, compression, and shear data.

## Discussion

The constitutive behavior of the human brain is highly complex, nonlinear, tension-compression asymmetric, and heterogeneous [6, 27]. Recent studies have shown that the classical principal-stretch-based Ogden model [15], initially developed for rubber, characterizes these features better than traditional invariant-based models [28]. However, because of its exponential nature, the general Ogden model is cumbersome to fit to experimental data [5]. Instead, most practical applications use only a limited subset of the Ogden model by either fixing its exponent [22], the number of terms [17], or both [21]. The full potential of Ogden-type modeling remains yet to be explored. Constitutive artificial neural networks [11] are a powerful new technology to integrate existing constitutive models into a neural network. As such, they harness the power of modern gradient-based optimizers that can robustly handle thousands of parameters [12]. We have recently prototyped this approach for an invariant-based neural network [14] and demonstrated that the network can autonomously discover the best constitutive model and parameters from a wide variety of existing models and combinations thereof [29]. Here we ask whether we can expand the concept of constitutive artificial neural network modeling to principal-stretch-based models and, if so, how these these networks perform compared to invariant-based networks.

### Principal-stretch-based constitutive artificial neural networks outperform invariant-based ones for human brain

After several preliminary studies, we selected a principal-stretch-based neural network with 20 Ogden-type terms [15] and exponents ranging from −30 to +10 in increments of two. Networks with more Ogden-type terms, a wider range of exponents, or narrower increments activated similar terms as the 20-term Ogden network and did not significantly alter our findings. When trained on all three loading modes, tension, compression, and shear [4], and averaged across the four brain regions, the principal-stretch-based network more than doubles the goodness of fit *R*^2^ for tension and improves the compression fit compared to the invariant-based network [14], bringing the R^2^ value close to the desired value of one for all loading modes. Additionally, for single-mode training in tension, compression, or shear, the principal-stretch-based network improves or nearly equals the fits across all loading modes compared to the invariant-based network. This agrees well with previous studies on brain tissue, in which the principal-stretch-based hyperfoam and Ogden models outperformed the invariant-based polynomial model [28]. In further support of Ogden type models, a study on brain and fat tissues found that only principal-stretch-based models were able to capture the characteristic behavior under combined stretch and shear, while classical invariant-based models failed to accurately capture the effects of combined loading [21].

### Low exponent Ogden-type models are not flexible enough to capture the pronounced tension-compression asymmetry in brain tissue

Popular constitutive models for fitting brain tissue data include the neo Hooke [23], Blatz Ko [24], and Mooney Rivlin [25, 26] models. The neo Hooke model uses a one-term Ogden model with an exponent of +2, the Blatz Ko uses a one-term model with exponent −2, and the Mooney Rivlin uses a two-term model with exponents −2 and +2. For single-mode training, compared to our twenty-term Ogden model, these low exponent models perform poorly at predicting tension behavior from compression data and vice versa. For multi-mode training with all three loading modes used in the training set, these low exponent models do not provide adequate fits in tension and are slightly worse in compression compared to the full principal-stretch-based constitutive artificial neural network. This poor performance becomes clear when comparing our twenty activation functions in Figure 2, where positive exponents *α* are associated with a stiffer behavior in tension than in compression and therefore not adequate for brain tissue. Negative exponents *α* are generally associated with a stiffer behavior in compression than in tension, but for the small stretch ranges, 0.9 ≤ *λ* ≤ 1.1, used in this study, large *α* values are necessary to translate into sufficient tension-compression asymmetry. This explains why our method consistently discovers red-colored terms, in Figures 4, 5, 6, and 8, which are associated large negative exponents *α*. These observations agree well with the initial analysis of the brain data [4], in which a principal-stretch-based one-term Ogden model with negative exponents in the range of *α* =-19 to −25 outperformed the classical neo Hooke, Mooney Rivlin, Demiray, and Gent models [6].

### Combined tension, compression, and shear data provide the most informative training data

Similarly to previous results from invariant-based neural networks [12], having all three loading modes in the training dataset for the principal-stretch-based network allows for the best model performance across all modes. Interestingly, the principal-stretch-based network without L1 regularization provides nearly as good a fit across all loading modes with just training on shear data. So, if we can only do a single experiment, for example because we want to limit post mortem time or have only limited availability of human brain tissue [6], our results indicate that shear is the most informative. Of course, if we desire the highest accuracy across all loading modes, training simultaneously on tension, compression, and shear data [4] is generally the best option.

## Conclusion

Principal-stretch-based models hold great promise to explain and predict the non-linear, tension-compression asymmetric behavior of soft biological tissues. However, the most prominent principal-stretch based model, the Ogden model, is cumbersome to calibrate because of its exponential nature. As a result, most existing Ogden-type models only represent a very small subset of possible models by a priori fixing the exponent, the number of terms, or both. Yet, the full potential of Ogden-type models remains underexplored. Here we propose a novel strategy to investigate principal-stretch-based constitutive modeling and embed the model into a constitutive artificial neural network to simultaneously discover both the relevant exponential terms and parameters that best describe experimental data. Rather than using a classical optimization approach, we harness the power of robust and efficient gradient-based adaptive optimizers developed for deep learning and train and test the network on human brain data. Our new principal-stretch-based network shows significant improvement over a recent invariant-based network at simultaneously training and testing on tension, compression, and shear data of gray and white matter tissue. When supplemented with L1 regularization, it tends to generalize better across all brain regions than the non-regularized network. Compared to other classical one-term models like the neo Hooke, Blatz Ko, and Mooney Rivlin models, our principal-stretch based network is able to discover the best model and parameters out of a possible combination of more than a million models, making it orders of magnitude more flexible to characterize the complex nonlinear, tension-compression asymmetric, ultrasoft behavior of human brain tissue. Taken together, our new principal-stretch-based constitutive artificial neural network autonomously discovers an optimal subclass of Ogden terms, their best parameters, and the best experiments for human brain tissue, making it a powerful tool for soft tissue modeling.

## Data Availability

Our source code, data, and examples are available at https://github.com/LivingMatterLab/CANN.

## Acknowledgments

This work was supported by a NSF Graduate Research Fellowship to Sarah St. Pierre, a DAAD Fellowship to Kevin Linka, and by the Stanford School of Engineering Covid-19 Research and Assistance Fund and Stanford Bio-X IIP seed grant to Ellen Kuhl.

## Notes

https://github.com/LivingMatterLab/CANN

### Competing Interest Statement

The authors have declared no competing interest.

## References

[1] Goriely, A. et al. Mechanics of the brain: Perspectives, challenges, and opportunities. Biomechanics Modeling and Mechanobiology 14, 931–965 (2015).

[2] Holzapfel, G. Nonlinear Solid Mechanics: A Continuum Approach for Engineering (Wiley, 2000).

[3] Zhao, W., Choate, B. & Ji, S. Material properties of the brain in injury-relevant conditions– experiments and computational modeling. Journal of the Mechanical Behavior of Biomedical Materials 80, 222–234 (2018).

[4] Budday, S. et al. Mechanical characterization of human brain tissue. Acta Biomaterialia 48, 319–340 (2017).

[5] Mihai, L. A., Budday, S., Holzapfel, G. A., Kuhl, E. & Goriely, A. A family of hyperelastic models for human brain tissue. Journal of the Mechanics and Physics of Solids 106, 60–79 (2017).

[6] Budday, S., Ovaert, T. C., Holzapfel, G. A., Steinmann, P. & Kuhl, E. Fifty shades of brain: a review on the mechanical testing and modeling of brain tissue. Archives of Computational Methods in Engineering 27, 1187–1230 (2020).

[7] Rashid, B., Destrade, M. & Gilchrist, M. D. Mechanical characterization of brain tissue in tension at dynamic strain rates. Journal of the Mechanical Behavior of Biomedical Materials 33, 43–54 (2014).

[8] Rashid, B., Destrade, M. & Gilchrist, M. D. Mechanical characterization of brain tissue in tension at dynamic strain rates. Journal of the Mechanical Behavior of Biomedical Materials 10, 23–38 (2012).

[9] Rashid, B., Destrade, M. & Gilchrist, M. D. Mechanical characterization of brain tissue in simple shear at dynamic strain rates. Journal of the Mechanical Behavior of Biomedical Materials 28, 71–85 (2013).

[10] Budday, S. et al. Mechanical properties of gray and white matter brain tissue by indentation. Journal of the Mechanical Behavior of Biomedical Materials 46, 318–330 (2015).

[11] Linka, K. et al. Constitutive artificial neural networks: A fast and general approach to predictive data-driven constitutive modeling by deep learning. Journal of Computational Physics 429, 110010 (2021).

[12] Linka, K. & Kuhl, E. A new family of constitutive artificial neural networks towards automated model discovery. Computer Methods in Applied Mechanics and Engineering 403, 115731 (2023).

[13] As’ad, F., Avery, P. & Farhat, C. A mechanics-informed artificial neural network approach in data-driven constitutive modeling. International Journal for Numerical Methods in Engineering 123, 2738–2759 (2022).

[14] Linka, K., St. Pierre, S. R. & Kuhl, E. Automated model discovery for human brain using constitutive artificial neural networks. bioRxiv doi:10.1101/2022.11.08.515656. (2022).

[15] Ogden, R. W. Large deformation isotropic elasticity–on the correlation of theory and experiment for incompressible rubberlike solids. Proceedings of the Royal Society of London. A. Mathematical and Physical Sciences 326, 565–584 (1972).

[16] Ogden, R. W. Non-Linear Elastic Deformations (SEllis Harwood Ltd., Chichester, Englandg, 1984).

[17] Lohr, M. J., Sugerman, G. P., Kakaletsis, S., Lejeune, E. & Rausch, M. K. An introduction to the ogden model in biomechanics: benefits, implementation tools and limitations. Philosophical Transactions of the Royal Society A 380, 20210365 (2022).

[18] Miller, K. & Chinzei, K. Mechanical properties of brain tissue in tension. Journal of Biomechanics 35, 483–490 (2002).

[19] Prange, M. T. & Margulies, S. S. Regional, directional, and age-dependent properties of the brain undergoing large deformation. Journal of Biomechanical Engineering 124, 244–252 (2002).

[20] Franceschini, G., Bigoni, D., Regitnig, P. & Holzapfel, G. A. Brain tissue deforms similarly to filled elastomers and follows consolidation theory. Journal of the Mechanics and Physics of Solids 54, 2592–2620 (2006).

[21] Mihai, L. A., Chin, L., Janmey, P. A. & Goriely, A. A comparison of hyperelastic constitutive models applicable to brain and fat tissues. Journal of The Royal Society Interface 12, 20150486 (2015).

[22] Ogden, R. W., Saccomandi, G. & Sgura, I. Fitting hyperelastic models to experimental data. Computational Mechanics 34, 484–502 (2004).

[23] Treloar, L. Stresses and birefringence in rubber subjected to general homogeneous strain. Proceedings of the Physical Society (1926-1948) 60, 135 (1948).

[24] Blatz, P. J. & Ko, W. L. Application of finite elastic theory to the deformation of rubbery materials. Transactions of the Society of Rheology 6, 223–252 (1962).

[25] Mooney, M. A theory of large elastic deformation. Journal of Applied Physics 11, 582–592 (1940).

[26] Rivlin, R. S. Large elastic deformations of isotropic materials iv. further developments of the general theory. Philosophical Transactions of the Royal Society of London. Series A, Mathematical and Physical Sciences 241, 379–397 (1948).

[27] Weickenmeier, J. et al. Brain stiffness increases with myelin content. Acta Biomaterialia 42, 265–272 (2016).

[28] Moran, R., Smith, J. H. & García, J. J. Fitted hyperelastic parameters for human brain tissue from reported tension, compression, and shear tests. Journal of Biomechanics 47, 3762–3766 (2014).

[29] Linka, K., Buganza Tepole, A., Holzapfel, G. A. & Kuhl, E. Automated model discovery for skin: Discovering the best model, data, and experiment. bioRxiv doi:10.1101/2022.12.19.520979. (2022).

